# Peripheral vs. Core Body Temperature as Hypocretin/Orexin Neurons Degenerate: Exercise Mitigates Increased Heat Loss

**DOI:** 10.1101/2022.12.21.521081

**Authors:** Yu Sun, Ryan K. Tisdale, Akira Yamashita, Thomas S. Kilduff

**Affiliations:** Biosciences Division, SRI International, 333 Ravenswood Ave, Menlo Park, CA 94025; Department of Physiology, Kagoshima University Graduate School of Medical and Dental Science, Kagoshima, 890-8544, Japan

**Keywords:** Hypocretin, Orexin, narcolepsy, body temperature, activity, thermogenesis, sympathetic vasoconstriction

## Abstract

Hypocretins/Orexins (Hcrt/Ox) are hypothalamic neuropeptides implicated in diverse functions, including body temperature regulation through modulation of sympathetic vasoconstrictor tone. In the current study, we measured subcutaneous (T_sc_) and core (T_b_) body temperature as well as activity in a conditional transgenic mouse strain that allows the inducible ablation of Hcrt/Ox-containing neurons by removal of doxycycline (DOX) from their diet (*orexin-DTA* mice). Measurements were made during a baseline, when mice were being maintained on food containing DOX, and over 42 days while the mice were fed normal chow which resulted in Hcrt/Ox neuron degeneration. The home cages of the *orexin-DTA* mice were equipped with running wheels that were either locked or unlocked. In the presence of a locked running wheel, T_sc_ progressively decreased on days 28 and 42 in the DOX(-) condition, primarily during the dark phase (the major active period for rodents). This nocturnal reduction in T_sc_ was mitigated when mice had access to unlocked running wheels. In contrast to T_sc_, T_b_ was largely maintained until day 42 in the DOX(-) condition even when the running wheel was locked. Acute changes in both T_sc_ and T_b_ were observed preceding, during, and following cataplexy. Our results suggest that ablation of Hcrt/Ox-containing neurons results in elevated heat loss, likely through reduced sympathetic vasoconstrictor tone, and that exercise may have some therapeutic benefit to patients with narcolepsy, a disorder caused by Hcrt/Ox deficiency. Acute changes in body temperature may facilitate prediction of cataplexy onset and lead to interventions to mitigate its occurrence.

**Highlights:**

- Hypocretin/Orexin (Hcrt/Ox) neuron degeneration results in the sleep disorder Narcolepsy and reduced subcutaneous body temperature (T_sc_) during the dark phase of the 24-h light/dark cycle.
- This reduction in dark phase T_sc_ is mitigated by access to an exercise opportunity.
- In contrast to T_sc_, core body temperature (T_b_) is largely maintained as the Hcrt/Ox neurons degenerate.
- Reduced T_sc_ while T_b_ is maintained suggests increased heat loss, possibly through modulation of sympathetic vasoconstrictor tone.
- Hcrt/Ox neuron loss in Narcolepsy results in cataplexy, whose occurrence is associated with acute changes in both T_sc_ and T_b_.
- Exercise may represent an effective intervention for mitigating heat loss resulting from Hcrt/Ox neuron loss in Narcolepsy.

## 1. Introduction

Hypocretins/Orexins (Hcrt/Ox) are hypothalamic neuropeptides implicated in diverse processes including sleep/wake control, addiction, the stress response through modulation of the hypothalamic-pituitary-adrenal (HPA) axis, as well as autonomic and body temperature regulation [1]. The two neuropeptides, hypocretin 1/orexin-A (Hcrt 1/Ox-A) and hypocretin 2/orexin-B (Hcrt 2/Ox-B), are both cleaved from the prepro-hypocretin/prepro-orexin protein, and have differential affinities for the Hypocretin 1/Orexin 1 (HcrtR1/OxR1) and Hypocretin 2/Orexin 2 receptors (HcrtR2/OxR2) [2, 3]. Hcrt/Ox-containing neurons project broadly and HcrtR/OxRs are distributed widely In the brain, thereby enabling this diverse functionality [4–8].

### 1.1. The role of Hcrt/Ox-containing neurons in body temperature and autonomic regulation

Although the link between the Hcrt/Ox system and metabolism was unequivocally established by the description of the *orexin/ataxin-3* mouse in which Hcrt/Ox neurons are ablated through selective expression of Ataxin-3 in Hcrt/Ox-containing neurons as becoming obese as the Hcrt/Ox neurons degenerate [9], this seminal paper did not describe effects of Hcrt/Ox neuron loss on body temperature regulation. Nonetheless, several lines of evidence have suggested a role for the Hcrt/Ox system in thermoregulation. For example, intracerebroventricular (ICV) administration of Hcrt1/Ox-A increased thermogenesis, leading to an elevation in body temperature in Sprague-Dawley rats [10]. Oral administration of a dual orexin receptor antagonist (DORA) resulted in reduced spontaneous locomotion for 8 hours and reduced core body temperature (T_b_) for 4 hours in Wistar rats [11]. Cold exposure increased *Hcrt/Ox* mRNA expression in the hypothalamus in Wistar rats [12]. Hcrt/Ox-containing neurons innervate neurons in the caudal raphe that project to medullary sympathetic premotor neurons that control brown adipose tissue (BAT) sympathetic outflow and thermogenesis [13–15]. In rats, intrathecal administration of Hcrt1/Ox-A increases sympathetic outflow [16]. Furthermore, local Hcrt1/Ox-A injection into the rostral raphe pallidus in rats induced prolonged and pronounced increases in BAT sympathetic outflow and thermogenesis [17]. Together, these studies are suggestive of a role for Hcrt/Ox neurotransmission in maintenance of body temperature through modulation of BAT sympathetic tone.

Because of the impact of the Hcrt/Ox system on a broad range of physiological functions including arousal state control, the precise role of this system in other autonomic processes has been difficult to elucidate [18]. Transgenic mice in which Hcrt/Ox neurotransmission (Hcrt/Ox-KO mice) is eliminated exhibit more frequent occurrence of central apneas during sleep and a reduction in CO_2_ response during wakefulness [19–21]. Furthermore, the ventilatory long term facilitation response is absent during both sleep and wake in Hcrt/Ox-KO mice [22]. Hcrt/Ox-KO mice and *orexin/ataxin-3* mice exhibit a basal blood pressure that is 20 mm Hg lower than their wildtype (WT) littermates [23–26]. Pharmacological blockade of sympathetic outflow lowered blood pressure in WT littermates and reduced basal blood pressure to levels similar to that found in Hcrt/Ox-KO mice [25]. The same pharmacological intervention had no effect in Hcrt/OX-KO mice, again suggesting reduced sympathetic vasoconstrictor tone as a result of the absence of the Hcrt/Ox neuropeptides, thus suggesting a causal role for impaired sympathetic tone in the reduction in basal blood pressure in the Hcrt/Ox-KO mice [25]. This role for the Hcrt/Ox system in maintenance of sympathetic vasoconstrictor tone thus has impacts on other autonomic processes such as modulation of cardiorespiratory function in addition to body temperature regulation.

### 1.2. Narcolepsy, a disorder caused by reduced or absent Hcrt/Ox neurotransmission: Metabolic disfunction and body temperature changes

Narcolepsy is a sleep disorder resulting from loss of Hcrt/Ox-containing neurons [27, 28], likely through an immune-mediated mechanism [29, 30]. The most prominent symptoms in Narcolepsy are related to sleep/wake and are a result of the loss of Hcrt/Ox input to wake-promoting regions such as the cholinergic basal forebrain neurons, resulting in reduced arousal state boundary control [30–32]. Patients with narcolepsy exhibit various sleep abnormalities including disrupted nocturnal sleep, sleep onset rapid-eye movement (REM) sleep, excessive daytime sleepiness (EDS) and cataplexy in Narcolepsy Type 1 (NT1) patients. People with NT1 also exhibit metabolic symptoms: they are more likely to be obese, are often hypophagic, and have a lower basal metabolic rate (BMR) than BMI-matched controls [33, 34]. Patients with NT 1 also display elevated plasma levels of GLP-1, suggestive of autonomic dysfunction [35].

Alterations in body temperature have also been described in patients with narcolepsy [36–38]. Increased skin temperature is associated with higher sleep propensity in NT1 [36]. An elevated dorsal-proximal skin temperature gradient (DPG) caused by elevated distal and reduced proximal skin temperatures during wakefulness is correlated positively with shorter sleep latency during a multiple sleep latency test (MSLT). During normal sleep, distal temperature remained elevated, while proximal skin temperature was similar to healthy individuals [36]. Altered DPG in patients with narcolepsy is suggestive of a decreased sympathetic vasoconstrictor tone. Hcrt/Ox-deficient patients with narcolepsy also exhibit a reduced heart rate response to arousal compared to healthy controls, lending further support to the concept of reduced BAT outflow occurring as a result of Hcrt/Ox deficiency [39]. This concept is further supported by the animal studies discussed in section 1.1 demonstrating that HcrtR1/OxR1agonism increases sympathetic outflow, suggesting that loss of Hcrt/Ox neurotransmission as occurs in narcolepsy would therefore result in decreased sympathetic output [16, 17].

In the current study, we utilized bigenic *orexin-tTA; TetO-DTA* mice, a strain in which ablation of Hcrt/Ox neurons can be induced through dietary manipulation [40]. Subcutaneous (T_sc_) and core (T_b_) body temperature were recorded in intact mice and over the course of a 42 days period as the Hcrt/Ox neurons degenerated. Since access to a running wheel has been reported to exacerbate cataplexy in orexin ligand knockout mice [41], we also investigated how provision of a running wheel and the consequent opportunity for exercise affected activity and body temperature.

## 2. Materials and Methods

### 2.1. Subjects

Male *“orexin-DTA* mice” used for EEG/EMG experiments were the double transgenic offspring of *orexin/tTA* mice (C57BL/6-Tg(*hOX-tTA*)1Ahky), which express the tetracycline transactivator (tTA) exclusively in Hcrt/Ox neurons [40], and B6.Cg-Tg(*tetO-DTA*)1Gfi/J mice (JAX #008468), which express a diphtheria toxin A (DTA) fragment in the absence of dietary doxycycline. Both parental strains were from a C57BL/6J genetic background. Parental strains and offspring used for EEG/EMG recordings were maintained on a diet (Envigo T-7012, 200 Doxycycline) containing doxycycline (DOX(+) condition) to repress transgene expression until neurodegeneration was desired. To initiate neurodegeneration, *orexin-DTA* mice were 15±1 weeks switched to normal chow (DOX(-) condition) to induce expression of the DTA transgene specifically in the Hcrt/Ox neurons [40]. All experimental procedures were approved by the Institutional Animal Care and Use Committee at SRI International and were conducted in accordance with the principles set forth in the *Guide for Care and Use of Laboratory Animals*.

### 2.2. Surgical Procedures

Male *orexin-DTA* mice (n = 31) were 11±1 weeks (23 ± 2 g) at the time of surgery. Mice were anesthetized with isoflurane (Induction: 3-5% isoflurane in oxygen delivered at 1 L/min; Maintenance: 1-2 % isoflurane in oxygen delivered at 1 L/min) and sterile telemetry transmitters (HD-X02, DSI, St Paul, MN) were placed subcutaneously (n = 26) for measurement of subcutaneous body temperature (T_sc_) or intraperitoneally (n = 5) for measurement of core body temperature (T_b_). Biopotential leads were routed subcutaneously to the head, and both EMG leads were positioned through the right nuchal muscles. Cranial holes were drilled through the skull at −2.0 mm AP from bregma and 2.0 mm ML and on the midline at −1 mm AP from lambda. The two biopotential leads used as EEG electrodes were inserted into these holes and affixed to the skull with dental acrylic. The incision was closed with absorbable suture. Analgesia was managed with meloxicam (5 mg/kg, s.c.) and buprenorphine (0.05 mg/kg, s.c.) upon emergence from anesthesia and for the first day post-surgery. Meloxicam (5 mg/kg, s.c., q.d.) was continued for 2 d post-surgery. Mice were monitored daily for 14 days post-surgery; any remaining suture material was removed at that time.

### 2.3. EEG, EMG, T_sc_, T_b_ and gross motor activity recording

Prior to data collection, *orexin-DTA* mice were allowed at least 2 weeks post-surgical recovery and at least 2 weeks adaptation to running wheels and handling procedures. Mice were housed individually in home cages equipped with running wheels and had *ad libitum* access to food, water and nestlets. Room temperature (22 ± 2°C), humidity (50 ± 20% relative humidity), and lighting conditions (LD12:12; Lights on at 7 am and off at 7 pm) were monitored continuously. Animals were inspected daily in accordance with AAALAC and SRI guidelines. EEG, EMG, T_sc_, T_b_ and gross motor activity (GMA) were recorded via telemetry using Ponemah (DSI, St Paul, MN). EEG and EMG were sampled at 500 Hz. Digital videos were recorded at 10 frames per second, 4CIF de-interlacing resolution.

### 2.4. Experimental Design

Two weeks after adaptation to running wheels, DTA mice were divided into 5 experimental groups (Table 1):

<u>Group 1</u>: *Orexin-DTA* mice implanted with telemetry transmitters subcutaneously to measure T_sc_ and maintained on DOX(+) diet for 42 days in cages with running wheels locked (n = 5);
<u>Group 2</u>: *Orexin-DTA* mice implanted with telemetry transmitters subcutaneously to measure T_sc_ and maintained on normal chow (DOX(-) condition) for 42 days in cages with running wheels locked (n = 5);
<u>Group 3</u>: *Orexin-DTA* mice implanted with telemetry transmitters subcutaneously to measure T_sc_ and maintained on normal chow (DOX(-) condition) for 42 days in cages with running wheels unlocked (n = 7);
<u>Group 4</u>: *Orexin-DTA* mice implanted with telemetry transmitters abdominally to measure T_b_ and maintained on normal chow (DOX(-) condition) for 42 days in cages with running wheels unlocked (n = 5).
<u>Group 5</u>: *Orexin-DTA* mice implanted with telemetry transmitters subcutaneously to measure T_sc_ and maintained on DOX(-) chow for 42 days. Following this DOX(-) period, initially mice were provided a locked running wheel cage, then after a 24 hour baseline recording beginning at ZT12 on day 0, the running wheels were unlocked at ZT 12 on the following day and 24 hour recordings were made on days 1, 2, 7, and 14 after unlocking the running wheels (n = 9).

**Table 1.**
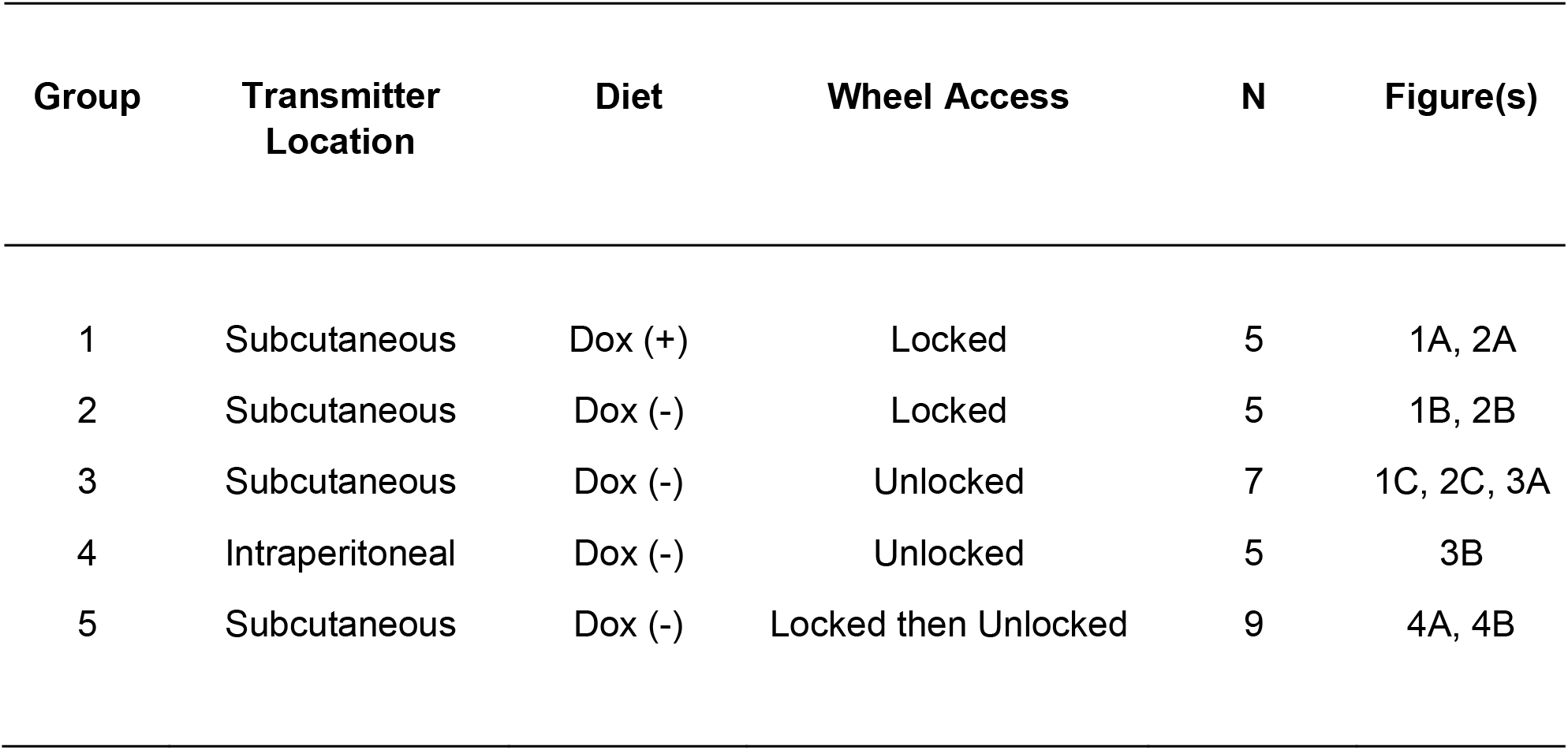
Description of experimental groups.

| **Group** | **Transmitter Location** | **Diet** | **Wheel Access** | **N** | **Figure(s)** |
| --- | --- | --- | --- | --- | --- |
| 1 | Subcutaneous | Dox (+) | Locked | 5 | 1A, 2A |
| 2 | Subcutaneous | Dox (-) | Locked | 5 | 1B, 2B |
| 3 | Subcutaneous | Dox (-) | Unlocked | 7 | 1C, 2C, 3A |
| 4 | Intraperitoneal | Dox (-) | Unlocked | 5 | 3B |
| 5 | Subcutaneous | Dox (-) | Locked then Unlocked | 9 | 4A, 4B |

For Groups 2-5, DOX(+) chow was reintroduced after 42 days to minimize further Hcrt/Ox neuron degeneration.

## 3. Results

### 3.1. Subcutaneous body temperature (T_sc_) and gross motor activity decline as the Hypocretin/Orexin neurons degenerate

Group 1 mice were maintained on the DOX(+) diet, resulting in suppression of transgene expression and, consequently, the Hcrt/Ox neurons should have remained intact. Thus, this group served as experimental controls and neither subcutaneous body temperature (T_sc_; Figs. 1A, A’, and A”) nor gross motor activity (Figs. 2A, A’, and A”) varied significantly from baseline over the 42 days observation period.

**Fig. 1.**
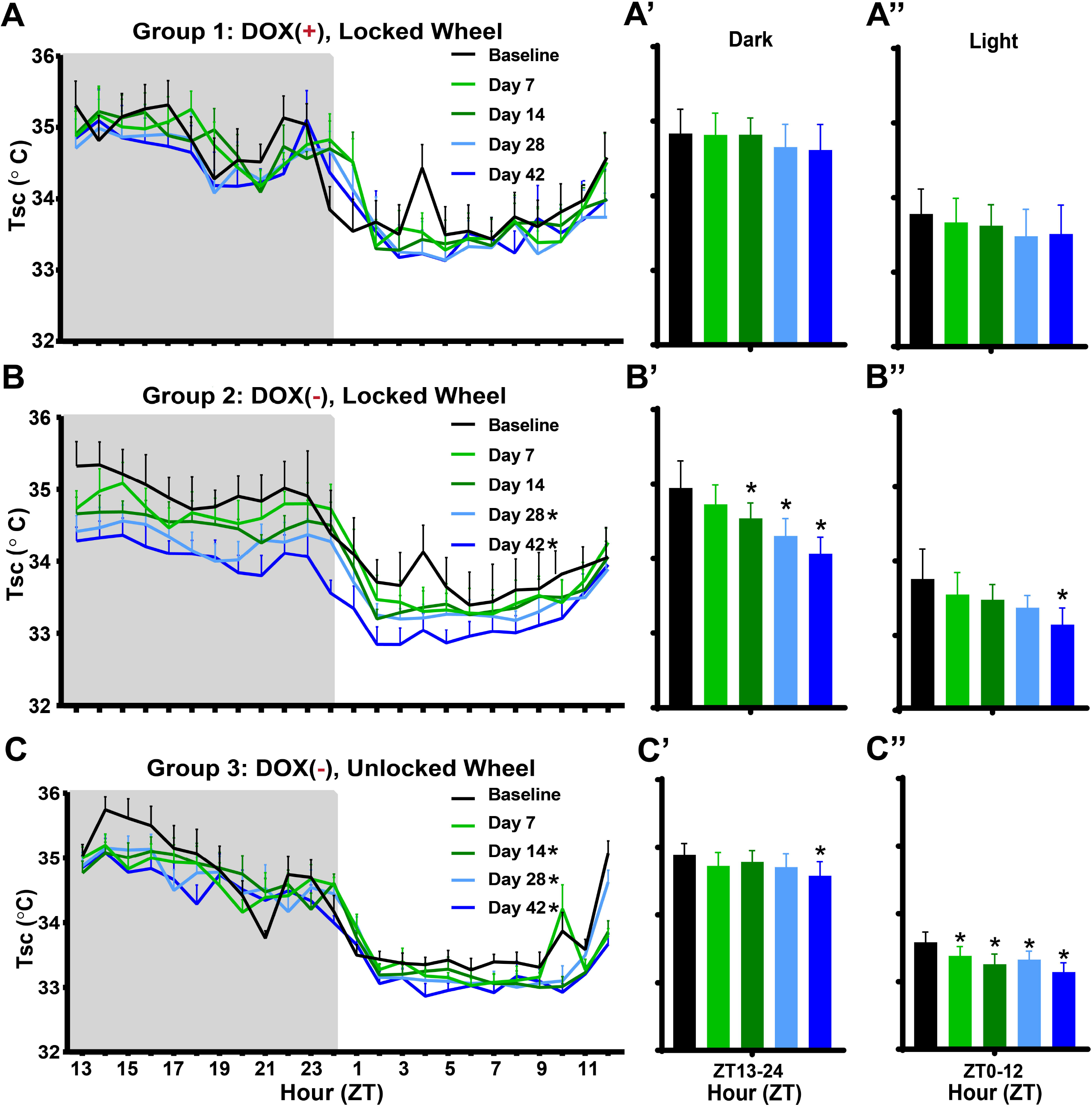
Effects of Hypocretin/Orexin neuron degeneration and access to a running wheel on subcutaneous body temperature (T_sc_) in male *orexin/tTA; TetO-DTA* mice. (A) Hourly T_sc_ across the 24-h period in Group 1 male *orexin/tTA; TetO-DTA* mice (n = 5) maintained on doxycycline (DOX(+)) with a locked running wheel in their home cage. Recordings were made during a baseline and 7, 14, 28, and 42 days later. (A’, A”) Mean T_sc_ over the 42 days period for the 12-h dark and light phases, respectively, for the mice recorded in A. (B) Hourly T_sc_ across the 24-h period in Group 2 male *orexin/tTA; TetO-DTA* mice (n = 5) that were maintained with a locked running wheel in their home cage. Mice were on DOX chow during baseline but then switched to normal chow (DOX(-) condition) for 42 days, during which time the Hcrt/Ox neurons degenerate. (B’, B”) Mean T_sc_ for the 12-h dark and light phases, respectively, during the 42-day degeneration period. (C) Hourly T_sc_ across the 24-h period in Group 3 male *orexin/tTA; TetO-DTA* mice (n = 7) that were maintained with an unlocked running wheel in their home cage. Mice were on DOX chow during baseline but then switched to normal chow (DOX(-) condition) for 42 days, during which time the Hcrt/Ox neurons degenerate. (C’, C’’) Mean T_sc_ for the 12-h dark and light phases, respectively, during the 42-day degeneration period. Values are mean ± SEM. * in the legend and above the vertical bars indicates a significant difference during that day relative to baseline as determined by ANOVA. * *p* < 0.05.

In contrast, when Group 2 mice were switched to normal chow (DOX(-) condition) after a baseline recording, T_sc_ decreased progressively over the course of Hcrt/Ox neuron degeneration during both the dark and light phases (Figs. 1B, B’, B’’). In comparison to the pre-degeneration baseline, RM-ANOVA revealed a significant condition effect for T_sc_ (*F*_(4, 16)_ = 6.467, *p* = 0.0027). Dunnett’s post-hoc tests indicate that T_sc_ was significantly reduced at days 28 (*p* = 0.0171) and 42 (*p* = 0.0009) after DOX removal. Fig. 1B’ shows that this reduction was most pronounced in the dark period on Days 14 (*p* = 0.0235), 28 (*p* = 0.0006), and 42 (*p* < 0.0001), as well as during the light period on day 42 (*p* < 0.0196) (Fig. 1B’’). A condition effect was also significant for gross motor activity (*F*_(4, 16)_ = 5.998, *p* = 0.0038). Gross motor activity (Fig. 2B) was significantly and progressively decreased at days 14 (*p* = 0.0378), 28 (*p* = 0.0026), and 42 (*p* = 0.0029). Figure 2B’ shows that this effect was primarily mediated by a reduction in gross motor activity during the dark period on days 14 (*p* = 0.01), 28 (*p* = 0.0012), and 42 (*p* = 0.0005), as activity didn’t vary significantly from baseline during the light period (Fig. 2B’’).

**Fig. 2.**
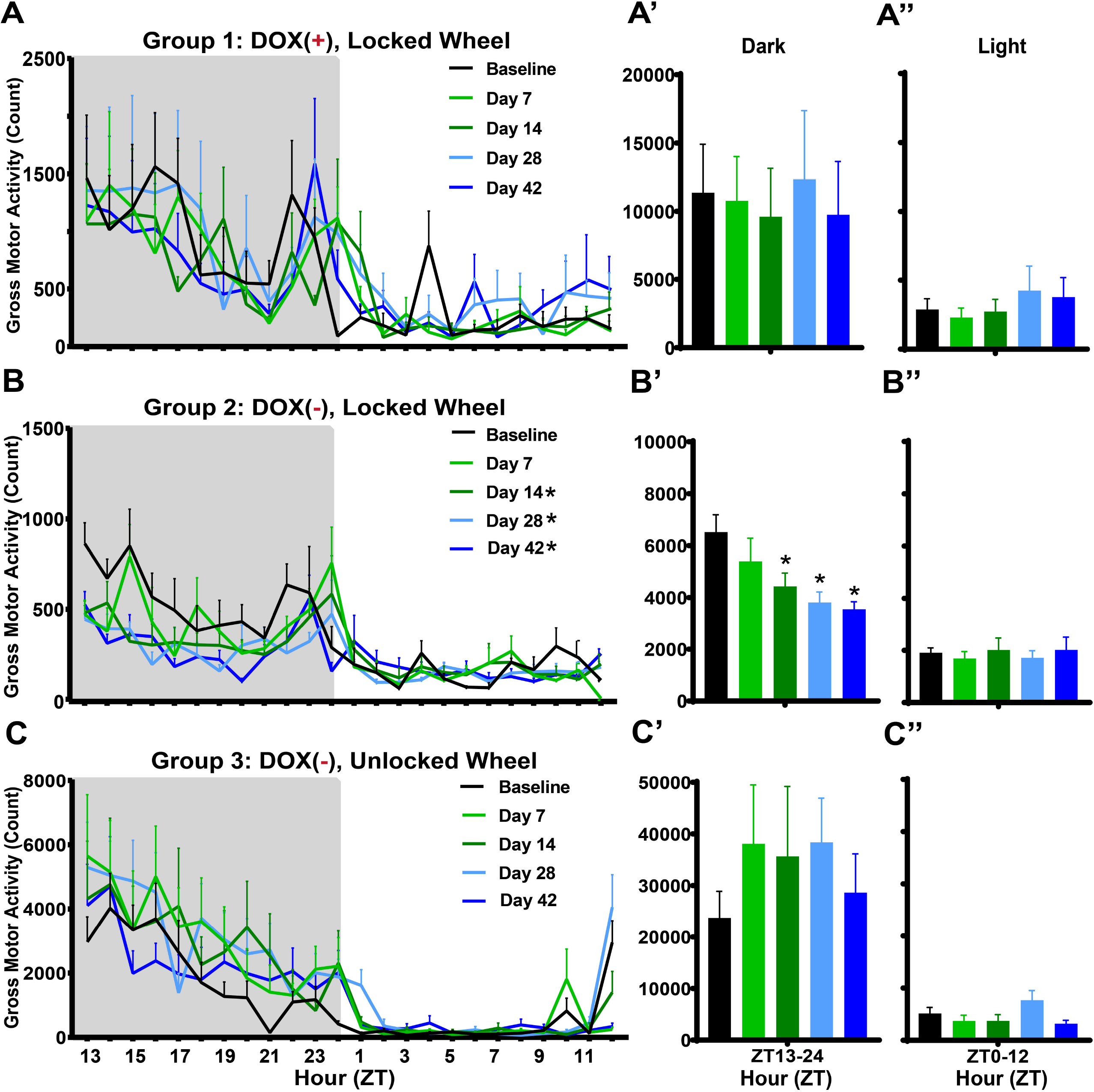
Effects of Hypocretin/Orexin neuron degeneration and access to a running wheel on Gross Motor Activity in male *orexin/tTA; TetO-DTA* mice. (A) Hourly Gross Motor Activity across the 24-h period in Group 1 male *orexin/tTA; TetO-DTA* mice (n = 5) maintained on doxycycline (DOX(+)) with a locked running wheel in their home cage. Recordings were made during a baseline and 7, 14, 28, and 42 days later. (A’, A”) Mean Gross Motor Activity over the 42-day period for the 12-h dark and light phases, respectively, for the mice recorded in A. (B) Hourly Gross Motor Activity across the 24-h period in Group 2 male *orexin/tTA; TetO-DTA* mice (n = 5) that were maintained with a locked running wheel in their home cage. Mice were on DOX chow during baseline and then switched to normal chow (DOX(-) condition) for 42 days, during which time the Hcrt/Ox neurons degenerate. (B’, B”) Mean Gross Motor Activity for the 12-h dark and light phases, respectively, during the 42-day degeneration period. (C) Hourly Gross Motor Activity across the 24-h period in Group 3 male *orexin/tTA; TetO-DTA* mice (n = 7) that were maintained with an unlocked running wheel in their home cage. Mice were on DOX chow during baseline and then switched to normal chow (DOX(-) condition) for 42 days, during which time the Hcrt/Ox neurons degenerate. (C’, C’’) Mean Gross Motor Activity for the 12-h dark and light phases, respectively, during the 42-day degeneration period. Values are mean ± SEM. * in the legend and above the vertical bars indicates a significant difference during that day relative to baseline as determined by ANOVA. * *p* < 0.05.

### 3.2. Access to a running wheel mitigates the decline in T_sc_ in the dark phase as Hypocretin/Orexin (Hcrt/Ox) neurons degenerate

In contrast to Groups 1 and 2, Group 3 mice had free access to a running wheel and their Hcrt/Ox neurons were degenerated by removal of dietary DOX as in Group 2 (Table 1). There was a significant condition effect on T_sc_ in Group 3 (*F*_(4,24)_ = 7.128, *p* = 0.0006). Similar to Group 2, T_sc_ was progressively reduced at days 14 (*p* = 0.0192), 28 (*p* = 0.0165), and 42 (*p* < 0.0001) post-DOX removal (Fig. 1C). In contrast to Group 2, this reduction was only significant at day 42 during the dark period (*p* = 0.0147; Fig. 1C’) but T_sc_ was significantly reduced during the light period at all post-DOX removal weeks in comparison to baseline (Day 7: *p* = 0.0226, Day 14: *p* = 0.0003, Day 28: *p* = 0.0031, Day 42: *p* < 0.0001; Fig. 1C”).

In contrast to Group 2 which experienced a reduction in gross motor activity as the Hcrt/Ox neurons degenerate (Figs. 2B, 2B’ and 2B”), Group 3 mice had similar activity levels across all recording conditions (Fig. 2C), both during the dark (Fig. 2C’) and the light (Fig. C”) phases.

### 3.3. T_sc_ declines as Hypocretin/Orexin (Hcrt/Ox) neurons degenerate even when a running wheel is available but core body temperature (T_b_) is maintained

Fig. 3 compares Groups 3 and 4 in which both groups of mice had free access to a running wheel while the Hcrt/Ox neurons degenerated due to removal of dietary DOX. However, in contrast to Group 3 in which T_sc_ was measured from a subcutaneously placed telemeter (Figs. 3A, 3A’ and 3A”; same data as Figs. 1C, 1C’ and 1C”), Group 4 mice had an intraperitoneally-placed telemeter which provided a measure of T_b_ (Figs. 3B, 3B’ and 3B”). While a condition effect was indicated on T_b_ in the group 4 (*F*_(4, 16)_ = 3.575, *p* = 0.0289), in distinction from Group 3 mice in which T_sc_ progressively declined from baseline on days 14 (*p* = 0.0192), 28 (*p* = 0.0165), and 42 (*p* < 0.0001) post-DOX removal (Fig. 3A), T_b_ was unchanged in Group 4 mice until Day 42 (*p* = 0.0066; Fig. 3B). T_b_ was most significantly altered during the dark period on day 42 post-DOX removal (*p* = 0.0198) in comparison to baseline (Fig. 3B’); no significant change in T_b_ was observed during the light phase (Fig. 3B’’).

**Fig. 3.**
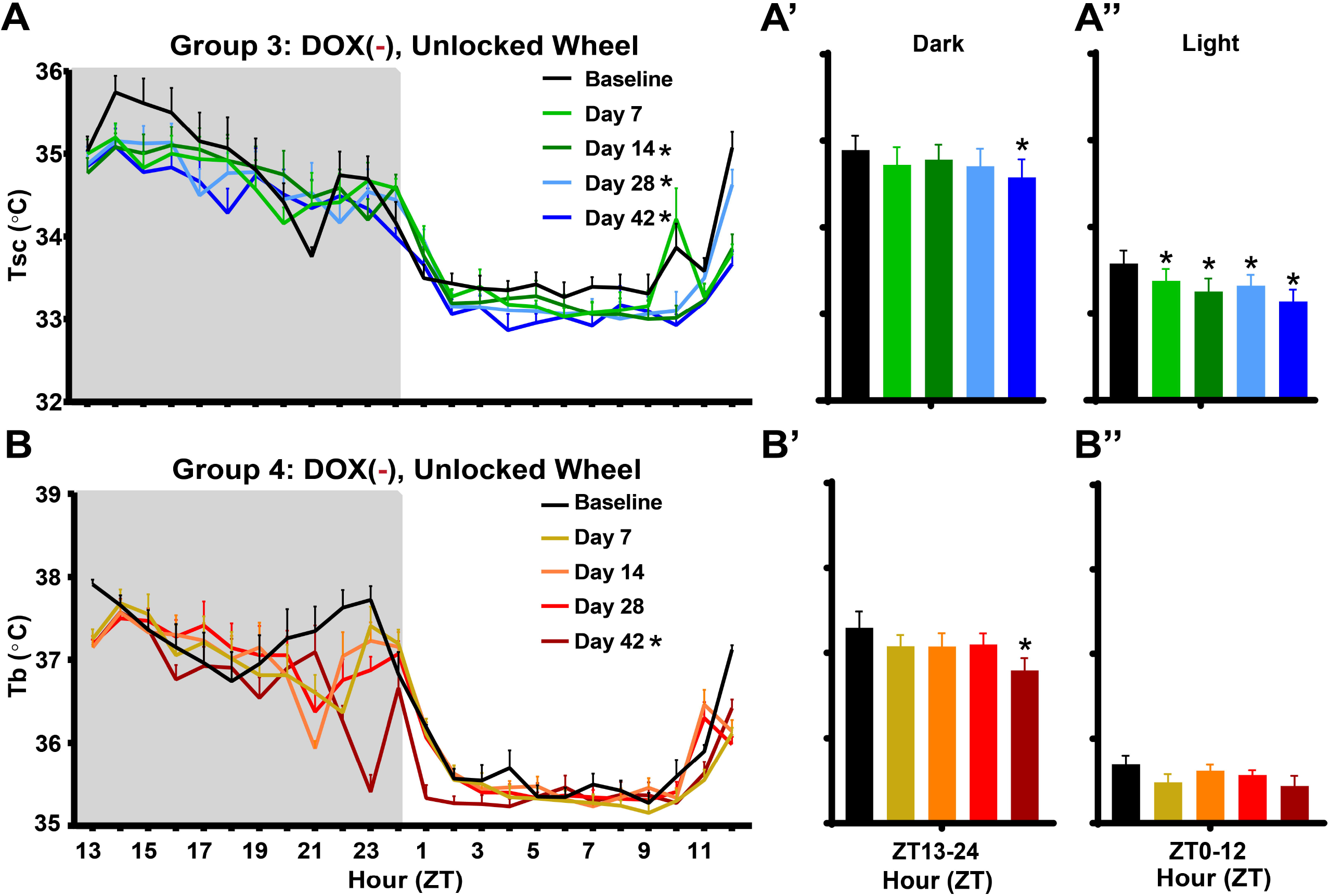
Peripheral vs. core body temperature as Hypocretin/Orexin neurons degenerate. (A, A’, A’’) Mean subcutaneous (T_sc_; n = 7) and (B, B’, B’’) core (T_b_; n = 5) body temperature in Group 3 vs. Group 4 male *orexin/tTA; TetO-DTA* mice initially maintained on DOX with an unlocked running wheel in their home cage. After a baseline recording, mice were switched to normal chow (DOX(-) condition) and recorded at 7, 14, 28, and 42 days after DOX removal during which time the Hcrt/Ox neurons degenerate. Values are mean ± SEM. * in the legend and above the bars indicates a significant difference during that day relative to baseline as determined by ANOVA. * *p* < 0.05.

Gross motor activity did not vary significantly over the course of Hcrt/Ox neuron degeneration in either Group 3 (Figs. 2C, 2C’ and 2C”), or Group 4 (data not shown).

### 3.4. Running wheel use in Hcrt/Ox neuron-degenerated mice elevates T_sc_, particularly during the dark phase

In Group 5, subcutaneously-implanted *orexin-DTA* mice were housed with a locked running wheel while Hcrt/Ox neuron degeneration proceeded in the DOX(-) condition for 42 days. After 42 days in the DOX(-) condition, a 24-hour recording of T_sc_ and gross motor activity was collected while the running wheel was still locked (Day 0). The wheel was then unlocked and recordings were collected on Days 1, 2, 7, and 14. Results from a two-way ANOVA indicated condition effects for both T_sc_ (*F*_(4, 32)_ = 4.046, *p* = 0.0091) and GMA (*F*_(4, 32)_ = 8.425, *p* < 0.0001). Post hoc tests indicated that after unlocking the wheel, T_sc_ and gross motor activity were indistinguishable on Days 1 and 2 from the locked condition. However, on Day 7 and Day 14, both T_sc_ (Day 7: *p* = 0.0024, and Day 14: *p* = 0.0412) and gross motor activity (*p* = 0.0003, and Day 14: *p* = 0.0002) were significantly increased compared to the locked condition (Day 0). This effect was most evident in the dark phase (T_sc_ Day 7: *p* = 0.0020, and T_sc_ Day 14: *p* = 0.0138; GMA Day 7: *p* = 0.0003, and GMA Day 14: *p* = 0.0002; Figs. 4A’ and 4B’). In the light period, T_sc_ (Fig. 4A’’) and gross motor activity (Fig. 4B’’) were only significantly increased on Day 7 (T_sc_: *p* = 0.0179 and GMA: *p* < 0.0001) compared to the locked condition (Day 0).

**Fig. 4.**
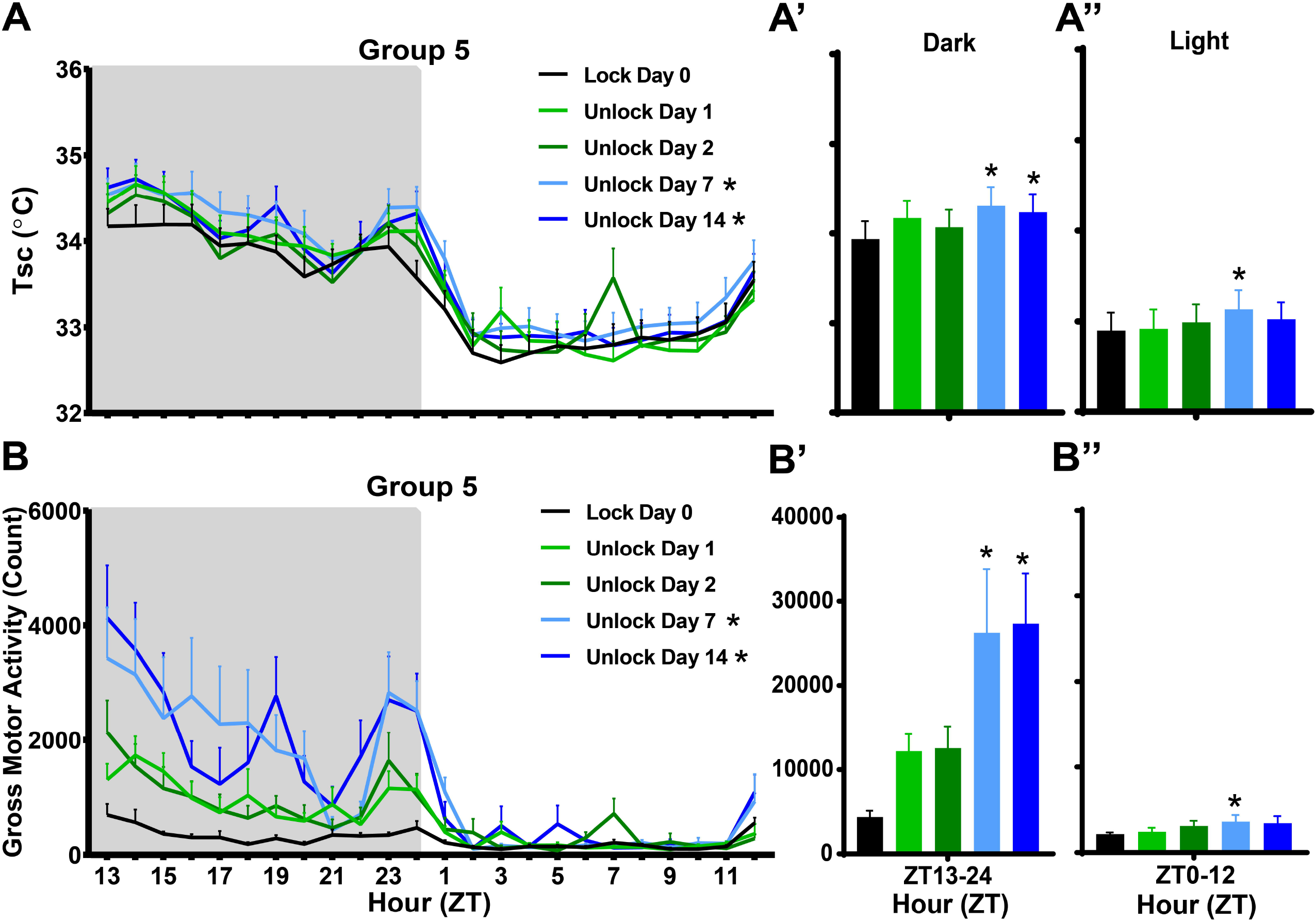
Peripheral body temperature and Gross Motor Activity changes as Hcrt/Ox neurondegenerated narcoleptic male *orexin/tTA; TetO-DTA* mice learn to use a running wheel. Mean subcutaneous (T_sc_; A, A’, A’’) body temperature and gross motor activity (B, B’, B’’) in Hcrt/Ox neuron-degenerated male *orexin/tTA; TetO-DTA* mice in Group 5 (n = 9) that were housed with a locked running wheel in their home cage. At the end of Day 0, the running wheel was unlocked just prior to light offset at ZT12 and recordings were conducted on days 1, 2, 7, and 14 after the wheel was unlocked. Values are mean ± SEM. * in the legend and above the bars indicates a significant difference during that day relative to the locked condition (Day 0) as determined by ANOVA. * *p* < 0.05.

### 3.5. Acute changes in T_sc_ and T_b_ during cataplexy bouts in Orexin-tTA;TetO-DTA mice

Removal of DOX from the diet of *orexin-DTA* mice produces Hcrt/Ox neurodegeneration and cataplexy occurs within 2 weeks in the DOX(-) condition [40, 42]. To the best of our knowledge, neither T_b_ nor T_sc_ had been systematically assessed during the period prior and subsequent to the onset of cataplexy. Figure 5 demonstrates that both T_sc_ and T_b_ changed acutely before, during, and after a bout of cataplexy. At 3 minutes before onset of a bout of cataplexy, T_sc_ is 35.39 ± 0.05°C and T_b_ is 36.27 ± 0.1 °C (Figs. 5A and B). Mean T_sc_ and T_b_ both increased prior to, and peak at, cataplexy onset. T_sc_ is significantly increased to 35.48 ± 0.06°C (*p* = 0.0229) and T_b_ to 36.36 ± 0.17°C (*p* < 0.0001) at cataplexy onset. During and after cataplexy, T_sc_ and T_b_ both decrease (Figs. 5A and B). At 3 minutes after cataplexy onset, T_sc_ is significantly decreased to 35.30 ± 0.06°C (*p* = 0.0001) and T_b_ to 36.28 ± 0.16°C (*p* = 0.0021).

**Fig. 5.**
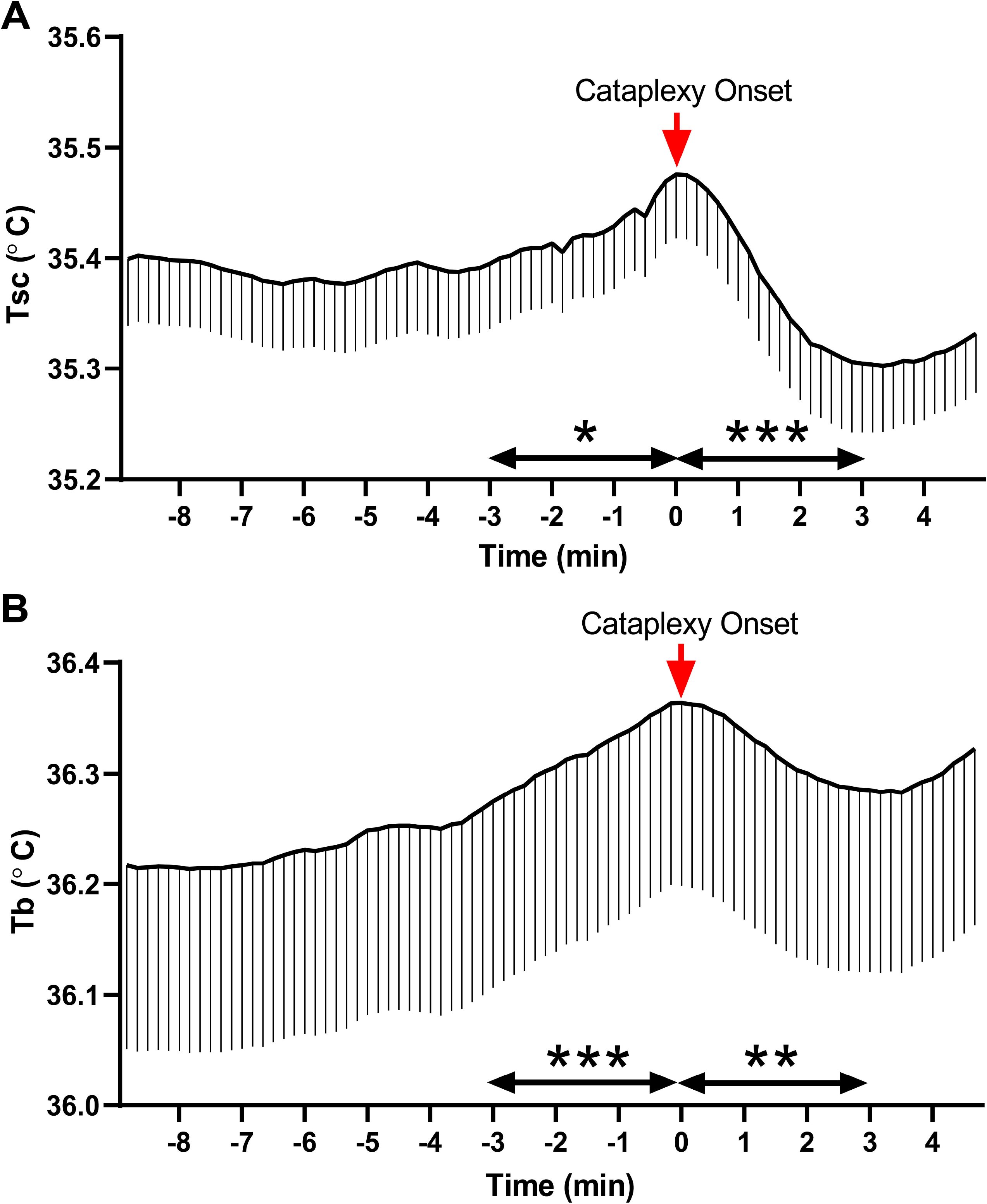
Acute effects of cataplexy occurrence on peripheral and core body temperature in male narcoleptic *orexin/tTA; TetO-DTA* mice. Subcutaneous (T_sc_) and core (T_b_) temperature change (mean ± SEM) from 9 min prior to cataplexy onset to 5 min after cataplexy onset. (A) T_sc_ determined from 33 cataplexy events from n = 7 mice. (B) T_b_ determined from 36 cataplexy events from n = 3 mice. Horizontal arrows indicate the T_sc_ and T_b_ change 3 min before and after cataplexy onset; vertical arrow (Red) indicates time of cataplexy onset. All mice were from the unlocked running wheel condition. ** *p* < 0.01, *** *p* < 0.001 vs Cataplexy onset T_sc_ and T_b_.

## 4. Discussion

### 4.1. Hcrt/Ox-containing neurons modulate heat loss and thereby affect core body temperature

From the perspective of temperature, the body can be divided into two broad regions, the internal “core” consisting of the central nervous system and viscera, and the external “shell” consisting of the skin, subcutaneous tissue, and limbs [43]. While core temperature is actively maintained, varying little from a set range, shell tissue is sensitive to environmental conditions. Shell temperature can also be reflective of autonomic thermoregulatory responses [44, 45]. In our narcoleptic mouse model, T_sc_ follows a similar trend to proximal skin temperature in humans with narcolepsy [36]. During the dark period, the major active period for rodents, T_sc_ is reduced as is proximal skin temperature during wakefulness in people with narcolepsy (Figs. 1B, B’ and B’’). This observation lends further support to notion that loss of Hcrt/Ox neurons results in reduced sympathetic tone, leading to elevated heat loss.

While T_b_ is maintained during the early stages of Hcrt/Ox neuron degeneration, it is significantly reduced by Day 42, primarily during the rodent’s major active phase (Figs. 3B, B’ and B’’). While these results are dissimilar to those obtained in patients with narcolepsy, who exhibited normal T_b_ during the day but increased T_b_ early in the night, in the current study, body temperature was measured in *orexin-DTA* mice over a period of acute degeneration [38]. Diagnosis in humans with narcolepsy often occurs years or decades after Hcrt/Ox neuron degeneration occurs [32]. Patients with narcolepsy also often present with comorbid obesity-related, metabolic conditions like diabetes myelitis, which is associated with altered body temperature, or hypertension, which occurs in conjunction with altered sympathetic tone [34, 46, 47]. Therefore, discrepancies between the human condition and the narcoleptic phenotype observed in degenerated *orexin-DTA* mice may be due to longer term network reorganization that inevitably occurs in the interim between development of the condition and diagnosis, or by comorbid metabolic disorders. As such, our results provide insight into the acute effects of loss of Hcrt/Ox neurons, something that is rarely observed clinically in patients with narcolepsy due to the difficulty in diagnosing narcolepsy in its prodromal phase. The acute reduction in T_b_ observed in this study may be due to the progressive reduction in sympathetic tone/increase in heat loss occurring as the Hcrt/Ox neurons degenerate, with heat loss exceeding thermogenic capability by Day 42 of degeneration.

Alternatively, REM sleep duration has recently been shown to correlate with basal body temperature [48]. The cumulative amount of REM sleep is largely unaltered in *orexin-DTA* mice [42]. However, the expression of cataplexy which, like REM sleep, is characterized by a sharp decrease in body temperature, develops progressively as the Hcrt/Ox neurons degenerate in DTA mice [42]. From a thermoregulatory perspective, cataplexy could therefore be equated with REM sleep and the reduced T_sc_ on Days 14, 28, and 42 of Hcrt/Ox-neuron degeneration as well as reduced T_b_ on Day 42 of degeneration in the present study may be due to increased cataplexy expression; however, this notion requires further study.

### 4.2. Hcrt/Ox neuron degeneration alters activity levels

Body temperature is elevated by sustained exercise and correlates positively with activity and changes in vigilance states [49–52]. In the current study, access to an unlocked running wheel mitigated the reduction in T_sc_ (Figs. 1C, 1C’, 1C’’, 4A, 4A’ and 4A’’) that occurred over the 42-day Hcrt/Ox neuron degeneration period in comparison to mice provided a running wheel that was locked (Figs. 1B, 1B’ and 1B’’), thus preventing the wheel to be used for running. In the locked condition, this T_sc_ deficit was most prominent during the dark period (Fig. 1B’). In Group 3 mice who had access to an unlocked running wheel, on the other hand, the T_sc_ deficits were mostly restricted to the light period (Fig. 1C’’). While Group 2 mice (locked running wheel) also exhibited a reduction in gross motor activity (Figs. 2A, 2A’ and 2A’’), the Groups 3 and 4 mice (unlocked running wheels) retained activity levels that were nearly identical to baseline (Figs. 2C, 2C’ and 2C’’; data not shown for Group 4). Despite similar activity levels in the unlocked condition, T_sc_ is still ultimately reduced in both groups.

Together, these results suggest impacts to at least two separate processes. While the progressive decrease in activity over the DOX(-) period in the mice provided locked running wheels contributed to the reduction in T_sc_ that occurred as the Hcrt/Ox neurons degenerated, the reduction in activity itself may reflect alterations to other processes, rather than changes in thermoregulation per se. *Orexin-DTA* mice exhibit alterations to sleep/wake when Hcrt/Ox neurons are degenerated, including a reduced ability to sustain wakefulness during the dark period, reflective of the excessive daytime sleepiness (EDS) that is characteristic of narcolepsy in humans [42]. Excessive sleepiness likely also causes the mice to be less active. The

Hcrt/Ox system is also implicated in regulating food intake, the arterial baroceptive reflex, and cardiorespiratory function [53, 54]. Responses of the Hcrt/Ox system immediately precedes autonomic responses [55]. Thus, the reduction in activity occurring as a result of loss of Hcrt/Ox input in *orexin-DTA* mice may be a secondary consequence of a metabolic phenotype resulting from alterations in any of these processes. Ultimately, both T_sc_ and T_b_ are reduced by Day 42 of the DOX(-) period even when activity is maintained in the unlocked running wheel condition, suggesting that elevated heat loss is occurring through reduced sympathetic tone, or reduced thermogenic capability in these mice after Hcrt/Ox neuron degeneration.

It is also of interest to consider why activity was progressively reduced in the presence of a locked running wheel (Figs. 2B, 2B’ and 2B’’), but is maintained when the mice are provided an unlocked running wheel (Figs. 2C, C’ and C’’). Running wheel access increases cataplexy expression in narcoleptic mice, suggesting that the ability to exercise in a running wheel provides a positive emotional stimulus, which is known to promote cataplexy occurrence [41]. It is thus tempting to speculate about a motivational or emotional component driving reduced activity in the absence of positive emotional stimulation since Hcrt/Ox neurotransmission has been implicated in various motivated behaviors [56–61]. Narcolepsy carries a high risk for psychiatric symptoms; however, it is unclear if this comorbidity is due to shared pathophysiology [62]. The acute reduction in activity observed in the current study in the absence of a positive emotional stimulus as Hcrt/Ox neurons degenerate in *orexin-DTA* mice could provide a system to study emotional and motivational aspects of narcolepsy.

### 4.3. Exercise reduces heat loss occurring during Hcrt/Ox neuron degeneration: Therapeutic implications in Narcolepsy?

In addition to pharmacological interventions currently approved to treat symptoms in patients with narcolepsy, lifestyle changes are often suggested to aid in symptom management. A low carbohydrate, ketogenic diet has been shown to improve narcoleptic symptomology as indicated by a decrease in patient score on the Narcolepsy Symptom Status Questionnaire and reduction in daytime sleepiness [63]. Cardiopulmonary fitness has an inverse relationship with sleepiness and cataplexy occurrence, lending support to exercise as an effective non-pharmacological intervention [64]. Thermoneutral core warming attenuated the normal decline in psychomotor vigilance test response speed in patients with narcolepsy, while distal skin cooling increased the patient’s ability to maintain wakefulness [37]. These results support non-pharmacological intervention through exercise as well as manipulation of body temperature as effective methods to mitigate narcolepsy symptoms. Figs. 3 and 4 suggest a relationship between exercise and rescue of the reduction in subcutaneously-measured body temperature occurring during Hcrt/Ox neuron degeneration, and thus support the use of exercise for symptom management in narcolepsy. The reduction in heat loss provided through exercise could reduce sleepiness and improve other narcoleptic symptoms.

The acute increases in body temperature preceding cataplexy onset observed in this study (Figs. 5A and B) suggest that body temperature could be a useful predictor for symptom onset. While body temperature changes before and during cataplexy have not been described clinically, a pronounced increase in body temperature has been previously noted to precede sleep attacks in humans with narcolepsy [36, 38]. These rapid changes in body temperature preceding symptom onset could be exploited to apply an acute intervention to mitigate cataplexy attacks.

### 4.4. Roles of Hcrt/Ox and colocalized neurotransmitters

In the conditional mouse strain used in this study, Hcrt/Ox-containing neurons are degenerated. While this is reflective of the narcoleptic condition in humans, Hcrt/Ox neurons coexpress other neuropeptides also implicated in body temperature regulation. In particular, glutamate and dynorphin have previously been implicated in body temperature regulation and are co-expressed in Hcrt/Ox-containing neurons [65–67]. Intriguingly, while Hcrt/Ox neuron-ablated mice [15, 21] and rats [68] have an attenuated thermogenesis response to stress, Prostaglandin E_2_ (PGE_2_)-induced fever, and cold stress, Hcrt/Ox-KO mice exhibit normal stress-induced hyperthermia, PGE_2_-induced fever, and mount a normal defense against cold stress [15, 21, 68]. These observations suggest a role for Hcrt/Ox neurons but not the Hcrt/Ox neuropeptides in BAT thermogenesis and highlights the importance of an intact Hcrt/Ox system in normal thermoregulation, but also underscores the importance of other molecules coexpressed in Hcrt/Ox-containing neurons in thermogenesis. The activity of putative Hcrt/Ox neurons lacking the Hcrt/Ox peptides increased immediately before the onset of cataplexy attack but decrease during attacks [69]. Local administration of a glutamate receptor antagonist to the dorsomedial hypothalamus inhibits BAT activation by PGE_2_, suggesting glutamate may be the critical modulator of sympathetic BAT stimulation by neurons in this region including Hcrt/Ox-containing neurons [70]. While the above data suggests colocalized neurotransmitters, most notably glutamate, may play a more critical role in thermoregulation, the studies cited in the Introduction that describe the effects of systemic Hcrt/Ox agonism and antagonism, as well as locally in the rostral raphe pallidus, support the involvement of Hcrt/Ox neurotransmission specifically in maintenance of sympathetic tone and BAT thermogenesis [11, 13–15, 17, 71].

## 5. Conclusions

Our results suggest that Hcrt/Ox-containing neurons are critical regulators of body temperature and autonomic functions related to activity and exercise. More research is needed to differentiate the contribution of the Hcrt/Ox peptides from other colocalized neurotransmitters in these processes. Acute changes in body temperature may predict cataplexy onset in narcolepsy, which could be exploited for the application of an acute intervention to prevent cataplexy/sleep attacks. Finally, exercise partially rescues the increase in heat loss that occurs as Hcrt/Ox neurons degenerate, supporting the use of exercise and other metabolic interventions for the treatment of narcolepsy.

## Abbreviations

BAT: Brown adipose tissue thermogenesis
DORA: Dual Orexin Receptor Antagonist
DOX: Doxycycline, GMA-Gross Motor Activity
Hcrt/Ox: Hypocretin/Orexin
Hcrt 1/Ox-A: Hypocretin 1/Orexin-A
Hcrt 2/Ox-B: Hypocretin 2/Orexin-B
HcrtR/OxR: Hypocretin/Orexin Receptor
PGE2: Prostaglandin E_2_
REM sleep: rapid-eye movement sleep
T_b_: Core body temperature
T_sc_: Subcutaneous body temperature
DTA: Diphtheria toxin A
Hcrt/Ox-KO mice: Prepro-hypocretin/prepro-orexin protein knockout mice

## Authorship contributions

Y.S., A.Y. and T.S.K. conceived of the study; Y.S. and A.Y. collected the data; A.Y. and Y.S. analyzed the data; R.K.T., Y.S. and T.S.K. drafted the manuscript; all authors edited and approved the final version of the manuscript.

## Acknowledgements

Research supported by NIH R01 NS098813 and R01 NS103529 to T.S.K. and Uehara Memorial Foundation 202040082 to A.Y.

